# STAT6 mRNA and protein knockdown using multiple siRNA sequences inhibits proliferation and induces apoptosis of the human colon adenocarcinoma cell line, HT-29

**DOI:** 10.1101/462895

**Authors:** C. Salguero-Aranda, D. Sancho-Mensat, S. Sultan, A. Reginald, L. Chapman

## Abstract

The transcription factor STAT6 is strongly expressed in various tumours and is most highly expressed in malignant lymphomas and pancreatic, colorectal, prostate and breast cancers. STAT6 expression in colorectal cancer is associated with an increased malignancy, poor prognosis and poor survival rates. Colorectal cancer has an incidence of approximately 1,361,000 patients *per annum* worldwide and approximately 60% of those cancers show STAT6 expression. Techniques aimed at reducing or blocking STAT6 expression may be useful in treating colorectal cancers. Celixir’s four proprietary STAT6 specific small interfering RNA (siRNA) sequences were tested *in vitro* using the human colon adenocarcinoma cell line, HT-29. The four sequences were introduced individually and in combination into HT-29 cells at different concentrations (10 to 200 nM). Decreases in STAT6 mRNA and protein levels were analysed to confirm the transfection was successful. STAT6 knockdown effects were measured by analysing cell proliferation and apoptosis. Results showed that 100nM siRNA concentration was the most effective and all four individual sequences knocked-down STAT6 mRNA and protein by more than 50%. Although all individual sequences were capable of significantly inhibiting cell proliferation, STAT6.1 and STAT6.4 were the best. STAT6 silencing also significantly induced late and total apoptotic events. In conclusion, these results demonstrate that STAT6 siRNA sequences are capable of inhibiting the proliferation, and inducing late apoptosis, of HT-29 colon cancer cells and, in some instances, halving the number of cancer cells. These experiments will be repeated using xenografts of STAT6-expressing colon cancer cells in immunocompromised mice and the STAT6 siRNA sequences will be tested in other cancers in which STAT6 is expressed. The STAT6 siRNA sequences therefore represent a potential treatment for the most serious colorectal cancers and a wide variety of STAT6-expressing cancers.

## Introduction

Colorectal cancer (CRC) represents 10% of cancers worldwide, ranking second in women and third in men (1). The incidence of CRC is approximately 1.36 million patients *per annum* worldwide (1). It is, overall, the fourth most common cause of death by cancer globally and its incidence is rising every year. Although most cases are detected in Western countries, its incidence is also increasing in developing countries (1,2). Actual CRC treatments involve a multimodal approach based on tumour characteristics and patient-related factors. Most CRC patients with metastases are treated with a combination of chemotherapy and targeted biological drugs but, in many cases, this is only a palliative approach (3). Therefore, the development of new, targeted and universal drugs for the treatment of CRC is needed.

The Signal Transducer and Activator of Transcription (STAT) family is formed by seven different transcription factors (STATs 1-4, 5a, 5b and 6). These proteins are important mediators in cytokine-related signalling and regulate normal cell differentiation, growth and survival (4). However, several of the STAT genes may be considered to be oncogenes (5). For example, STAT3 is overexpressed and active in many types of cancer, and its targeting by specific inhibitors is being deeply investigated as a potential cancer treatment (6). STAT6 has also been implicated in cancer. STAT6 is principally activated by two cytokines in the physiologic setting: interleukin-4 and interleukin-13 (7–11). Once these cytokines bind to their cell surface receptors, associated Janus Kinases (Jak) are activated and phosphorylate tyrosine residues on the receptors. Cytoplasmic STAT6 docks onto the phosphorylated receptors allowing the Jaks to phosphorylate the conserved tyrosine-641 on STAT6. Once phosphorylated, two STAT6 proteins form a homodimer and the homodimer translocates to the nucleus where it can directly regulate transcription (9). STAT6 has a well-known role in tumour immunosurveillance, immune function and lymphomagenesis but has only recently been associated with cancer progression. The STAT6 pathway has been heavily studied in animal models. STAT6-defective mice have shown immunity to mammary carcinoma (12) and also spontaneous rejection of implanted tumours (13). In humans, high levels of STAT6 have been detected in different cancer types, including glioblastoma, lymphoma, colorectal, prostate, pancreatic, and breast cancer (14). In addition, different studies have shown how STAT6 signalling pathway activation may be involved in the development of prostate, breast and colon carcinoma (11, 15–17). Moreover, in CRC, STAT6 is associated with increased malignancy and poor prognosis, and patients with CRC expressing STAT6 also show poor survival rates (18). The 5-year relative survival rate for patients with stage IIIC and IV colon cancer is approximately 53% and 11% respectively (19). Therefore, techniques aimed at reducing STAT6 expression may be useful in treating those cancers.

Gene silencing by double-stranded (ds) RNA-mediated interference (RNAi) was first described by Craig Mello and his colleagues in 1998 (20), for which they were awarded the Nobel Prize in 2008. It is a simple and rapid method of silencing gene expression in a range of organisms by degradation of RNA into small interfering RNAs (siRNAs) that activate ribonucleases to target homologous messenger RNA (mRNA) (21). siRNAs occur naturally from different sources (repeat-associated transcripts, viral RNAs, hairpin RNAs, *etc*) but can also be synthesized chemically and introduced into the cells. siRNAs are formed by two strands: the guide strand that assembles into a functional siRNA RNA-induced silencing complex (siRISC), which binds to an Ago protein, and a passenger strand that is discarded and degraded. The siRISC complex recognizes target RNAs by base pairing with the guide strand, leading to the silencing of the target gene through one of several mechanisms (22). Due to its superb specificity and efficiency, siRNA is considered as an important tool for gene-specific therapeutic activities that target the mRNAs of disease-related genes.

Consequently, the development of nucleotide-based biopharmaceuticals is a flourishing industry. According to recent reviews, more than 14 siRNA therapeutics have entered clinical trials in the past decade (23).

In this study, the potential effects of four proprietary STAT6 siRNA sequences, previously tested for asthma treatment (24), in a colon cancer cell line were examined to test the hypothesis that knocking-down STAT6 can prevent the proliferation and survival of CRC cells.

## Material and Methods

### Cell culture

Human colon adenocarcinoma cell line HT-29 was acquired from the European Collection of Authenticated Cell Cultures (ECCAC) (Catalogue Number 91072201, ATCC^®^ HTB-38). HT-29 cells were cultured in McCoy’s 5a medium (Sigma Aldrich) supplemented with 10% fetal bovine serum (FBS) (Sigma Aldrich), 2 mM of L-Glutamine (Sigma Aldrich), 100 U/ml of penicillin and 100 μg/ml of streptomycin (Sigma Aldrich) at 37°C and 5% CO_2_. Cells were passaged when 80-90% confluence was reached, and the media was changed every 2 - 3 days.

### siRNA transfection

Cells were seeded in 6-well and 12-well plates at a concentration of 15,000 cells/cm^2^. 24 hours post-culture, cells were then transfected with the four siRNA sequences at different final concentrations using DharmaFECT Transfection Reagent 1 (Dharmacon) or jetPEI (Polyplus) in antibiotic-free media, following the manufacturer’s instructions. jetPEI transfection was developed using a ratio of reagent:siRNA of 2:1. The senses of the STAT6 siRNA sequences were: Sequence 1 (STAT6.1): 5′ GCAGGAAGAACUCAAGUUUUUUU 3′, Sequence 2 (STAT6.2): 5′ ACAGUACGUUACUAGCCUUUUUU 3′, Sequence 3 (STAT6.3): 5′ GAAUCAGUCAACGUGUU GUUUUU 3′, Sequence 4 (STAT6.4): 5′ AGCACUGGAGAAAUCAUCAUUUU 3′. Sequential transfections were developed using STAT6.1 and STAT6.4 at 100 nM. Non-targeting siRNA and GAPDH (Dharmacon) were used as negative and positive controls respectively at 10 to 200 nM, depending on the assay. Media was not changed until the first 48 hours and antibiotic-free media was always used.

### RNA isolation, reverse transcription and q-PCR

After 24 hours of transfection, total RNA was isolated using a microRNA Isolation Kit (Qiagen) according to the manufacturer’s instructions directly from the plate. mRNA was then quantified by Nanodrop 1000-ND and 1 μg of RNA was transcribed into complementary DNA (cDNA) using SuperScript III Reverse Transcriptase (Invitrogen) following the manufacturer’s instructions. The DNA primers used were Human STAT6 Forward: 5′ CTTTCCGGAGCCACTACAAG 3′ and reverse 5′ AGGAAGTGGTTGGTCCCTTT 3′; Human GAPDH Forward: 5′ - TGCACCACCAACTGCTTAGC 3′ and reverse 5 ′ GGCATGGACTGTGG TCATGAG 3′. The quantitative PCR (qPCR) was performed in a 7900HT Real-time PCR system (ThermoFisher). The program cycle was: initial denaturation for 5 min at 95°C, followed by 40 cycles of 15 sec at 95°C and 60 sec at 60°C. A melt curve was added at the end of the process. The data was analysed by Delta-Delta Ct method.

### STAT6 protein detection

48 hours post-transfection, cells were harvested and fixed and permeabilized with Cell Signalling Buffer Set A (Miltenyi) according to the manufacturer’s instructions. In brief, cells were fixed for 10 min at room temperature (RT) with the Inside Fix Buffer and permeabilized for 30 min at 4°C with the Permeabilization Buffer pre-cooled at −20°C. Cells were washed twice with PBS/0.5%BSA and stained with anti-STAT6 APC conjugated antibody (Miltenyi Biotec, 130-104-030) (20 μl/10^6^ cells) and anti-GAPDH FITC conjugated antibody (Millipore, 130-104-030) (2 μl/ 10^6^ cells) for 30 min in the dark at 4°C. Antibody isotypes REA Control (I)-FITC (Miltenyi, 130-104-611) and REA Control (I)-APC (Miltenyi, 130-104-615) were used as controls. The stained cells were washed once and resuspended finally in 400 μL of PBS/0.5%BSA, before analysing them by flow cytometry (FACSCalibur, BD). Data was analysed using FlowJo software (FlowJo, BD).

### Cell proliferation

Cells were grown for 3, 6 or 8 days to analyse individual transfection, and 13 and 15 days for sequential transfection assays. 48 hours post-transfection the media was replaced, and every 2 days, fresh antibiotic-free media was added. Cell number has been used as a measure for cell proliferation. Total and dead cells were counted using a NucleoCounter NC-100 (Chemometec) and live cells were then calculated.

### Apoptosis analysis

Cells were harvested 7 days after transfection. Cells where then stained with anti-Annexin V FITC-conjugated antibody (BD Bioscience, 556420) at 20 μl/1 × 10^6^ cells in Binding Buffer 1X (BD Bioscience), for 15 min RT protected from light. Cells were finally resuspended in 400 μl of Binding Buffer 1X and 100 μl of propidium iodide (PI) solution (250 nM) (Sigma-Aldrich) was added to the cells and incubated for 1 min before analysing with the flow cytometer (FACSCalibur, BD). Data was analysed using FlowJo software (FlowJo, BD).

### Statistical analysis

The statistical analysis was carried out using PRISM software, by Student’s t-distribution of unpaired data, two-tailed, and 95% level of confidence. Values were compared to the non-targeted condition. Significant values: *(p-value <0.05), **(p-value<0.01), ***(P<0.001), ****(P-value<0.0001).

## Results

### STAT6 siRNA optimal dose and best sequences

In order to test the four proprietary STAT6 siRNA sequences’ efficiency, the first step was to determine the optimal dose. Ascending concentrations of STAT6: 10, 25, 50, 100 and 200 nM were tested twice (2n) for each STAT6 siRNA sequence and both STAT6 mRNA and protein levels were measured. Results illustrated all four sequences worked efficiently at silencing STAT6 expression. All conditions tested showed significant changes versus cells treated with non-targeting (NT) siRNA, with the exception of 10 and 25 nM of STAT6 sequence 2 (STAT6.2) and 10 nM of STAT6 sequence 3 (STAT6.3) at mRNA level. Regarding the expression of the protein, all conditions showed statistically significant changes, with 100 and 200 nM being the most effective, achieving an average of more than 60% knockdown for the four sequences. No significant changes were observed between 100 and 200 nM (S1A and B Fig). For this reason, 100 nM was established as the STAT6 siRNA optimal dose and this concentration was used for the remaining assays. To determine the effects of STAT6 siRNA on HT-29 cell proliferation, cells were transfected with 100 nM of each siRNA sequence and counted at different time points. Results showed that STAT6.2 and STAT6.3 reduced the number of live cells after 8 days in culture by approximately 20-30%, while STAT6 siRNA sequences 1 (STAT6.1) and 4 (STAT6.1) achieved a reduction of approximately 50% (S2A and B Fig). The reduction of the total number of cells when STAT6.1 and STAT6.4 were used was also appreciable under the inverted microscope (S2C Fig). To demonstrate the efficacy of STAT6.1 and STAT6.4 when used at 100 nM, more biological replicates were developed and these clearly demonstrated that STAT6 expression was reduced by approximately 50% at both mRNA and protein level (Fig 1A and B). Flow cytometer analyses revealed that STAT6 fluorescence was extremely decreased in STAT6.1 and STAT6.4 transfected cells (Fig 1C and D). Thereby, 100 nM and STAT6.1 and STAT6.4 were established to be the optimal dose and best sequences respectively, and they were used for the subsequent experiments.

In order to test the four proprietary STAT6 siRNA sequences’ efficiency, the first step was to determine the optimal dose. Ascending concentrations of STAT6: 10, 25, 50, 100 and 200 nM were tested twice (2n) for each STAT6 siRNA sequence and both STAT6 mRNA and protein levels were measured. Results illustrated all four sequences worked efficiently at silencing STAT6 expression. All conditions tested showed significant changes versus cells treated with non-targeting (NT) siRNA, with the exception of 10 and 25 nM of STAT6 sequence 2 (STAT6.2) and 10 nM of STAT6 sequence 3 (STAT6.3) at mRNA level. Regarding the expression of the protein, all conditions showed statistically significant changes, with 100 and 200 nM being the most effective, achieving an average of more than 60% knockdown for the four sequences. No significant changes were observed between 100 and 200 nM (S1A and B Fig). For this reason, 100 nM was established as the STAT6 siRNA optimal dose and this concentration was used for the remaining assays. To determine the effects of STAT6 siRNA on HT-29 cell proliferation, cells were transfected with 100 nM of each siRNA sequence and counted at different time points. Results showed that STAT6.2 and STAT6.3 reduced the number of live cells after 8 days in culture by approximately 20-30%, while STAT6 siRNA sequences 1 (STAT6.1) and 4 (STAT6.1) achieved a reduction of approximately 50% (S2A and B Fig). The reduction of the total number of cells when STAT6.1 and STAT6.4 were used was also appreciable under the inverted microscope (S2C Fig). To demonstrate the efficacy of STAT6.1 and STAT6.4 when used at 100 nM, more biological replicates were developed and these clearly demonstrated that STAT6 expression was reduced by approximately 50% at both mRNA and protein level (Fig 1A and B). Flow cytometer analyses revealed that STAT6 fluorescence was extremely decreased in STAT6.1 and STAT6.4 transfected cells (Fig 1C and D). Thereby, 100 nM and STAT6.1 and STAT6.4 were established to be the optimal dose and best sequences respectively, and they were used for the subsequent experiments.

**Fig1.**
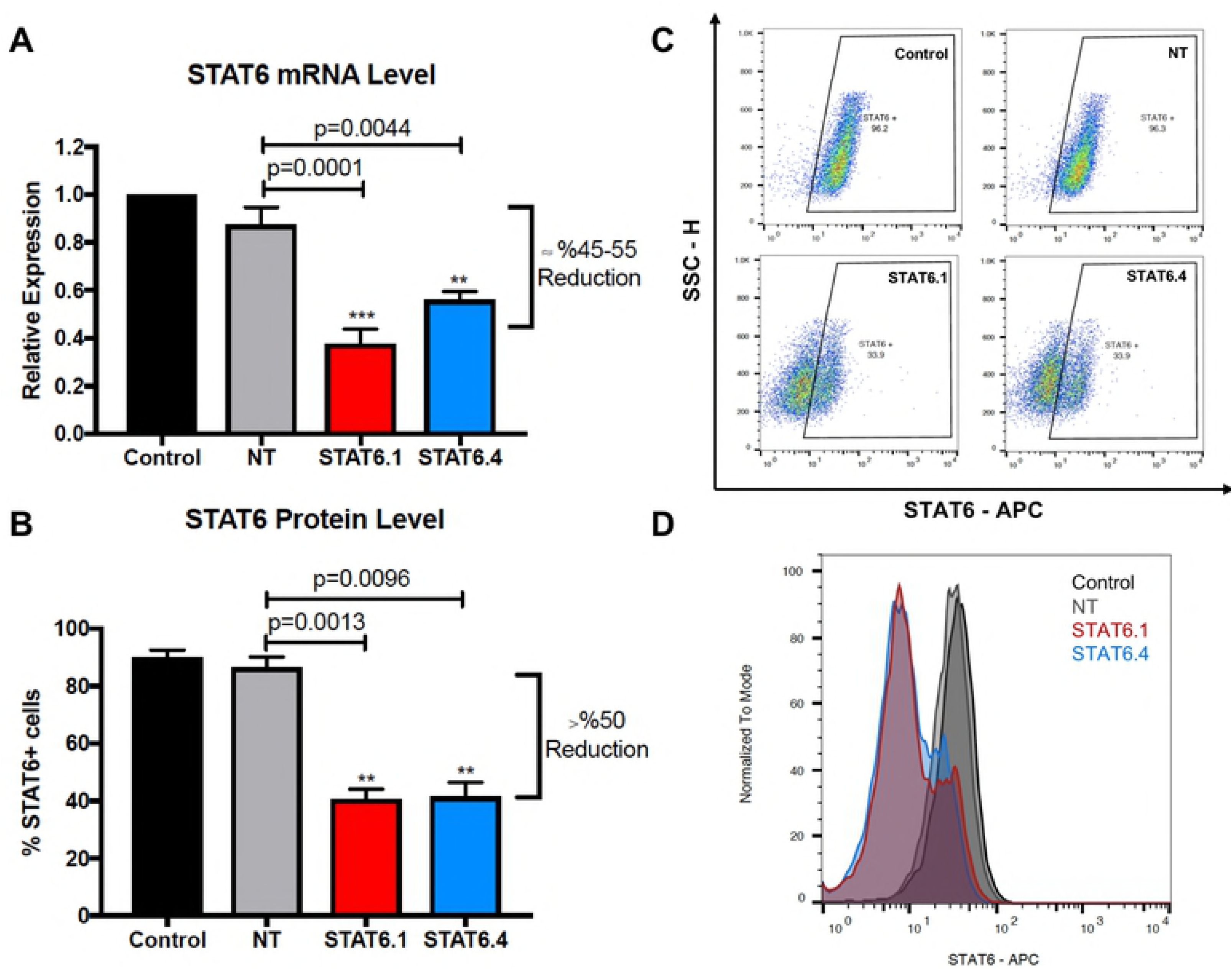
STAT6 siRNA sequences 1 and 4 (STAT6.1 and STAT6.4) powerfully block STAT6 expression. (A) STAT6 mRNA level measure. The graph represents the mean ± SEM of 6 (Control, NT and STAT6.1) or 3 (STAT6.4) independent experiments obtained by real-time PCR. Results were analysed by ∆∆Ct method for relative quantifications. The fold change is represented by the Y axis, and values are normalized to control cells. (B) STAT6 protein level analysis. The graph represents the mean of the percentage of STAT6 positive cells ± SEM of 6 (Control, NT and STAT6.1) or 5 (STAT6.4) independent experiments obtained by flow cytometry. (C) Representative dot plot and (D) histogram of STAT6 protein analysis by flow cytometry. STAT6 siRNA sequences and non-targeting siRNA were used at 100 nM as the final concentration. Control cells were non-transfected cells and STAT6 siRNA sequences 1 and 4 and non-targeting siRNA are denoted as STAT6.1, STAT6.4 and NT, respectively.

### STAT6 siRNA sequences 1 and 4 (STAT6.1 and STAT6.4) are highly efficient in silencing STAT6 expression

Employing the methods used previously, a number of biological replicates (STAT6.1, n=7 and STAT6.4, n=3) were analysed to analyse cell proliferation post-transfection with 100 nM of STAT6.1 and STAT6.4. NT cells had a similar growth pattern to the control cells, and STAT6.1 and STAT6.4 treatments significantly reduced HT-29 cell proliferation. At both 6 and 8 days of culture, approximately 50% of the number of live cells were obtained post-transfection with STAT6.1 and STAT6.4 in comparison with cells transfected with NT (Fig 2). Moreover, an increased concentration of STAT6.1 and STAT6.4 was tested (200 nM), but no significant changes were seen (data not shown). In addition, combinations of two, three and four STAT6 siRNA sequences 1, 2, 3 and 4 were also studied. However, there was no improvement in the results obtained (data not shown). These experiments demonstrate that the STAT6 siRNA sequences, and especially STAT6.1 and STAT6.4, are capable of significantly reducing the number of cancer cells *in vitro* in a short period of time.

**Fig2.**
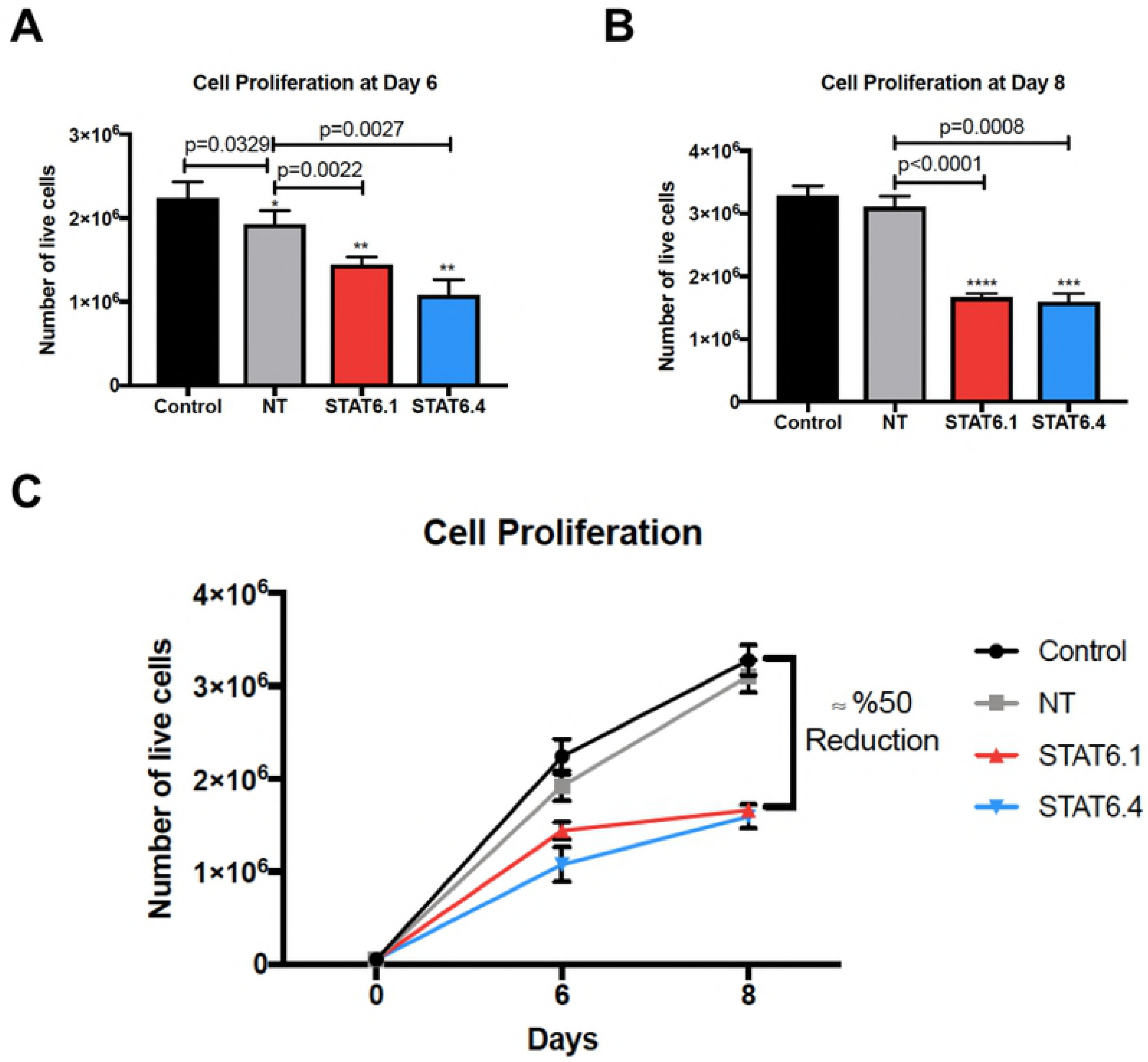
STAT6 siRNA sequences 1 and 4 (STAT6.1 and STAT6.4) significantly reduce cell proliferation. (A) Number of live cells measured at day 6 of culture. The graph represents the mean ± SEM of 7 (Control, NT and STAT6.1) or 4 (STAT6.4) independent experiments. (B) Number of live cells measured at day 8 of culture. The graph represents the mean ± SEM of 8 (Control, NT and STAT6.1) or 5 (STAT6.4) independent experiments. (C) The graph illustrates how cells grew over time and represents the mean ± SEM of the independent experiments shown in A and B. The number of live cells was calculated as detailed in the material and methods using NucleoCounter NC-100. STAT6 siRNA sequences and non-targeting (NT) siRNA were used at 100 nM as the final concentration. Non-transfected cells served as negative controls and STAT6 siRNA sequences 1 and 4 and non-targeting siRNA are denoted as STAT6.1, STAT6.4 and NT, respectively.

### STAT6 siRNA sequences induce apoptotic events

Once STAT6.1 and STAT6.4 were shown to significantly reduce the number of live cells over time, the implication that STAT6 also induces apoptosis was tested. After 8 days of culture, cells were harvested and counterstained with Annexin V and PI. Cells were then analysed by flow cytometry and the results showed that the percentage of Annexin V^+^/PI^+^ cells was increased in approximately 40% and approximately 50% of cells, when HT-29 cells were transfected with STAT6.1 and STAT6.4, respectively, compared to NT (Fig 3A and C). Furthermore, the number of total apoptotic events (Annexin V^+^ cells) was also augmented in both cases (Fig 3B and C). Moreover, 200 nM and a combination of STAT6 siRNA sequences were also tested and apoptosis was measured, however, an improvement in data was not observed (data not shown).

**Fig3.**
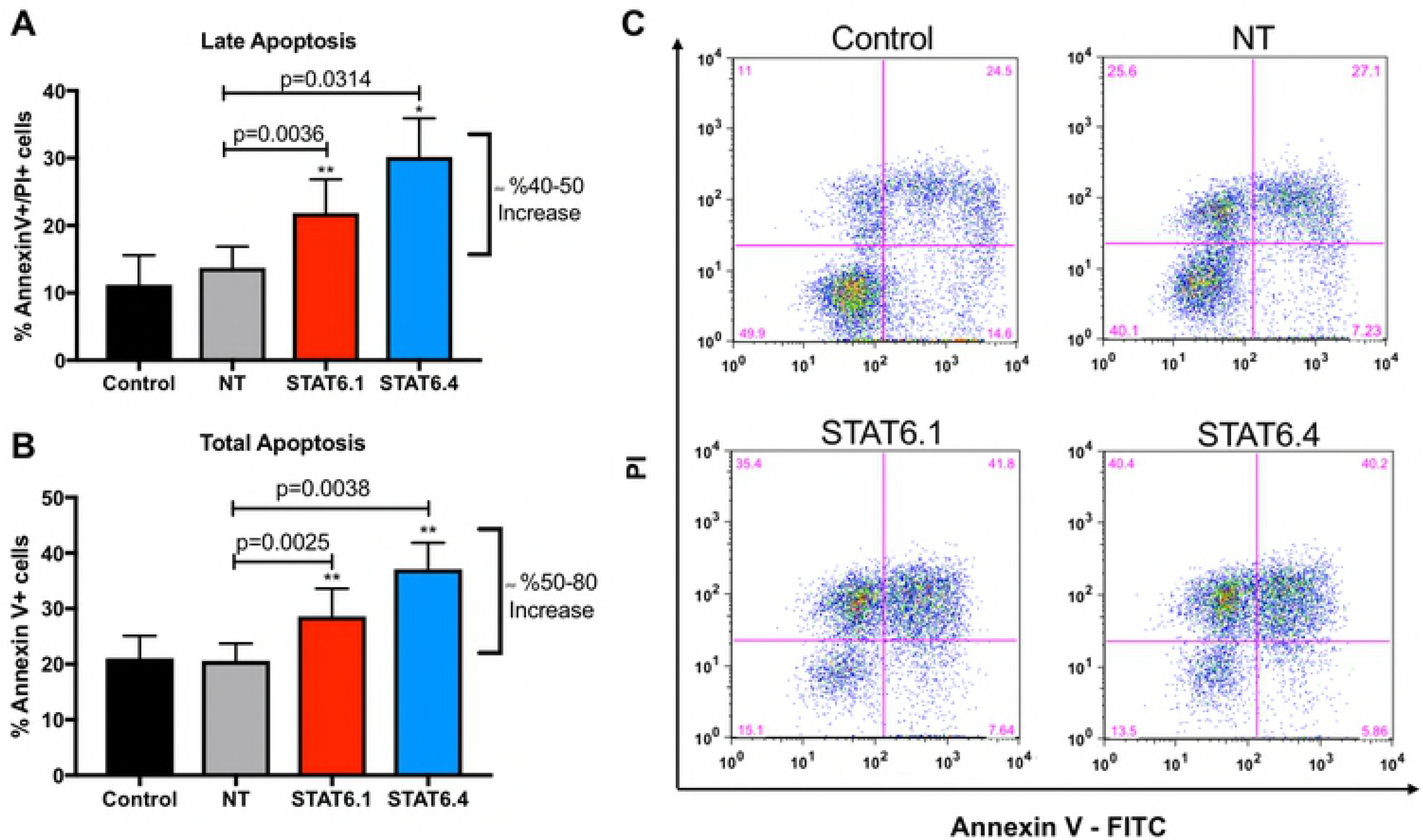
STAT6 siRNA sequences 1 and 4 (STAT6.1 and STAT6.4) induce apoptosis. (A) Late Apoptosis: percentage of Annexin V and PI positive cells. (B) Total Apoptosis: percentage of Annexin V positive cells. The graphs represent the mean ± SEM of 7 (Control, NT and STAT6.1) or 5 (STAT6.4) independent experiments obtained by flow cytometry. (C) Representative flow cytometry plots. The X axis represents Annexin V and the Y axis, PI fluorescence intensity. Quadrants were set according to cells independently stained with Annexin V or PI. Apoptosis was studied 7 days post-transfection and data were analysed with Flowjo Software. STAT6 siRNA sequences and a non-targeting siRNA sequence were used at 100 nM as the final concentration. Non-transfected cells served as control cells and STAT6 siRNA sequences 1 and 4 and non-targeting siRNA are denoted as STAT6.1, STAT6.4 and NT, respectively.

**Fig4.**
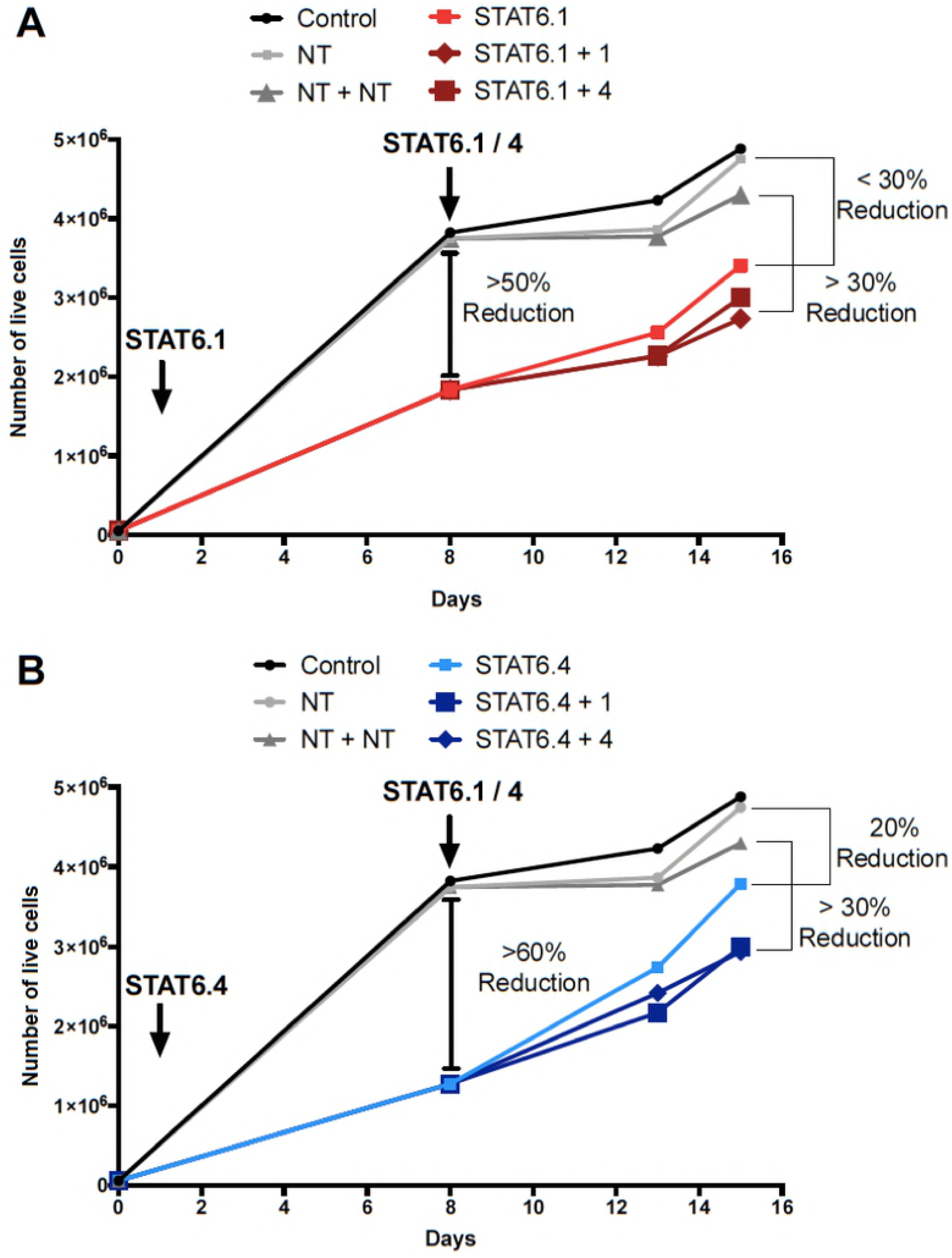
STAT6 siRNA serial transfection is effective in maintaining a reduced number of cells over time. STAT6 siRNA transfection was carried out at day 1 of cell culture with (A) STAT6.1 and (B) STAT6.4 at 100 nM. A second transfection was carried out in both cases with STAT6.1 and STAT6.4 at the same concentration 7 days after the first transfection. The graphs represent the number of live cells over time measured at day 8, 13 and 15 counted using NucleoCounter NC-100 as detailed in the material and methods section. The values were obtained from 1 independent experiment. Control cells were non-transfected cells and STAT6 siRNA sequences 1 and 4 and non-targeting siRNA are denoted as STAT6.1, STAT6.4 and NT, respectively. The percentage of reduction of the number of live cells was calculated by comparison between the mean of NT *vs*. the mean of STAT6 siRNA sequences individual transfection, and double transfection with NT (NT+NT) *vs*. double transfection with STAT6.1 and STAT6.4.

### STAT6 siRNA sequential transfection works at maintaining a reduced number of cancer cells over time

It is clear from these results that STAT6.1 and STAT6.4 at 100 nM can significantly reduce the number of live CRC cells cultured for up to 8 days. Further experiments were conducted to see if the effects of the siRNA sequences could be extended. Serial transfection using STAT6.1 and STAT6.4 at 100 nM each transfection was developed. First, transfection was prepared as usual, and 7 days later, a second transfection was performed using the same STAT6 siRNA sequences (STAT6.1 or STAT6.4) or the other STAT6 siRNA. The cells were cultured for a total of 15 days. The results showed that STAT6.1 and STAT6.4 individually achieved less than approximately 30% reduction of the number of live cells, while the serial combination achieved more than a 30% reduction, regardless of the combination used (Fig 5A and B). These data confirm that serial injections in animal models could be effective extending the effects of the siRNA sequences.

**Fig 5.**
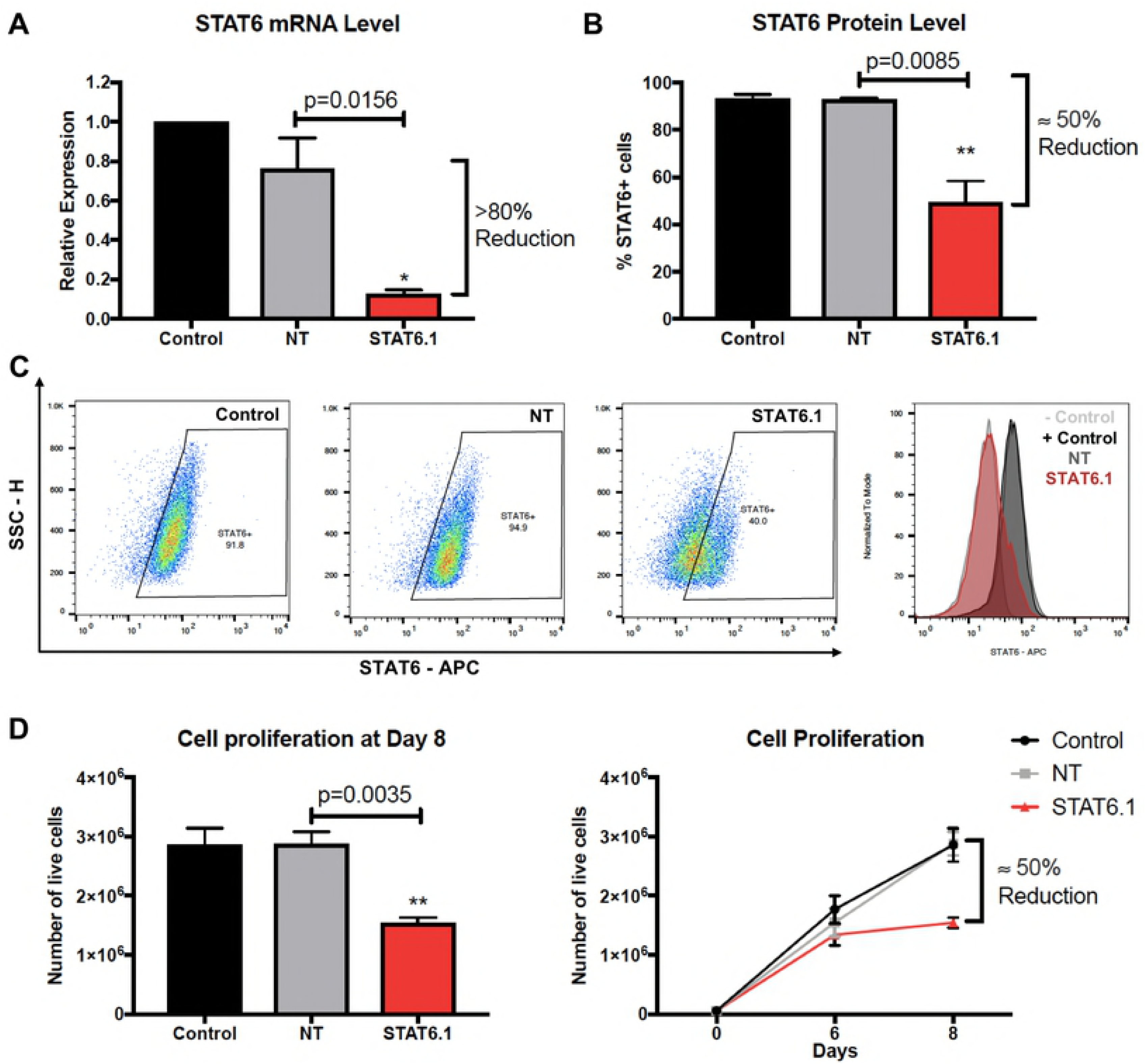
JetPEI transfection reagent works for STAT6 siRNA treatment in vitro. (A) STAT6 expression at mRNA level. The graph represents the mean ± SEM of 3 independent experiments. Total mRNA was measured by real-time PCR and results were analysed by the ∆∆Ct method for relative quantifications and values were normalized to control cells. (B) STAT6 expression at protein level. The graph represents the mean ± SEM of 3 independent experiments. Data was analysed using Flowjo Software for MacOS. The percentage of STAT6 positive cells is represented on the Y axis. (C) Representative dot plots and histogram from one set of experiments. STAT6 fluorescence is represented on the X axis. (D) Cell proliferation analysis. Number of live cells measured at day 6 and 8 of culture. The graphs represent the mean ± SEM of 3 independent experiments. The number of live cells was calculated as detailed in the material and methods section using NucleoCounter NC-100.

### JetPEI transfection reagent works for STAT6 siRNA treatment *in vitro*

The previous experiments were conducted using DharmaFECT, a lipid-based transfection reagent that provides efficient and reliable transfection at low concentrations with minimal cellular toxicity, but its use has not been tested *in vivo*. Therefore, once it was established that STAT6.1 and STAT6.4 individually at 100 nM had significant effects on cell proliferation and apoptosis of HT-29 cells, the efficacy of these STAT6 siRNA sequences was tested using a transfection reagent with proven efficacy *in vivo*. jetPEI reagent is a linear polyethylenimine derivative, free of components of animal origin, providing a highly effective and reproducible gene delivery to adherent and suspension cells and with a similar composition to *in vivo*-jetPEI, which is widely use in *in vivo* studies. In this case, only STAT6.1 was tested. Results using jetPEI for transfection showed again that STAT6.1 signifcantly silenced STAT6 expression, obtaining approximately 80% and approximately 50% knockdown at the mRNA and protein levels, respectively, compared with NT cells (Fig 5A and B). Fig 5C shows how STAT6 fluorescence was decreased in silenced HT-29 cells. The next step was to analyse if the effects of STAT6.1 on HT-29 cell proliferation and apoptosis were reproducible when jetPEI was used. The results showed that after 8 days of culture, the number of live cells were significantly decreased, obtaining approximately 50% reduction of the number of live cells (Fig 5D). However, no significant induction in apoptosis was observed (data not shown). These results show that the jetPEI transfection reagent could be an option for future animal studies.

## Discussion

CRC represents the fourth most common cause of death by cancer in the world and its incidence is increasing every year (1,2). Despite many efforts, the prognosis of CRC is still poor (19). Thus, exploring the underlying mechanism of CRC and finding new treatment targets are essential for improving the survival rate of CRC patients. Several studies have shown that STAT6 plays an important role in the progression and proliferation of several different types of cancer. Barbara C Merk *et al*. demonstrated in 2011 (25) that STAT6 acts to enhance cell proliferation and invasion in glioblastoma, which may explain why up-regulation of STAT6 correlates with shorter survival times in glioma patients. A study in 2007 showed that the actions of STAT6 in lung cancer were directly involved in COX-2 expression (26). A more recent study suggests that miR-135b functions as a tumour suppressor, affecting the metastatic ability of prostate cells by targeting STAT6, and STAT6 knockdown resulted in reduced cell metastasis. Furthermore, the expression of miR-135b was observed to be associated with the pathological T stages and levels of total and free PSA in patients with prostate cancer (27). It has been also shown that the inhibition of the STAT6 pathway in tumor-associated macrophages (TAMs) is a vital therapeutic approach to attenuate tumor growth and metastatic niche formation in breast cancer (28). In the same way, Yan D. *et al*. have determined that cytokine-activated STAT3 and STAT6 cooperate in macrophages to promote a secretory phenotype that enhances tumor progression in a cathepsin-dependent manner (29). STAT6 is also associated with an increased malignancy and a poor prognosis in CRC patients (18). Moreover, it has been demonstrated that the IL-13/IL-13Rα1/STAT6/ZEB1 pathway plays a critical role in promoting aggressiveness of CRC (30). It is for these reasons STAT6 was chosen in this study as a key target in CRC cells and the reported results suggest that the STAT6 siRNA sequences, especially STAT6.1 and STAT6.4, have the potential to treat CRC.

This study is not the first time that STAT6 knockdown in HT-29 has been investigated. Zhang MS *et al*. showed in 2006 that STAT6-specific short hairpin RNAs (shRNAs) inhibit proliferation and induce apoptosis in CRC HT-29 cells (31). They analysed the expression of total STAT6 and phosphorylated STAT6 protein by semiquantitative RT-PCR, obtaining a significant reduction of the STAT6 expression. HT-29 cell viability was also tested 72 hours post-transfection, and the results showed a greatly decreased viability. Apoptosis analysis by flow cytometry (Annexin V and PI) indicated that STAT6 shRNAs induced significant early apoptotic events (Annexin V^+^/ PI-cells). In this study, STAT6.1 and STAT6.4 also induced late apoptosis (Annexin V^+^/ PI^+^). This may be due to the fact that the apoptosis assay was analyzed after 7 days post-transfection, which would allow the STAT6 pathway to complete its action mechanism, or that the STAT6 siRNA sequences are more powerful at inducing the apoptosis of the cancer cells. In this study, the effects of STAT6 siRNA over a longer period of time (7 and 15 days) were investigated and this provided new data regarding the effects of STAT6 on cell proliferation and apoptosis. Moreover, Zhang *et al.* used shRNA, which is expressed after nuclear delivery of an shRNA-expressing plasmid DNA (pDNA), and the duration of shRNA expression depends on the use of viral or non-viral vectors. Conversely, the delivery of siRNAs as in this study avoids the barrier of the nuclear membrane as it acts in the cytosol (32). siRNAs offer additional advantages over shRNAs. Pre-designed siRNA duplexes are available from various sources or can be custom designed. Furthermore, siRNAs are easy to modify to increase their stability without altering their structure and efficiency and can be conjugated with fluorophores for *in vivo* tracking. In addition to this, the amount of exogenous nucleic acid introduced into the cells is much lower, as siRNAs consist of only duplexes of 19 nucleotide pairs and no insertion vector is required, thus reducing probable side effects.

It is for these and other reasons why siRNAs are becoming a popular tool for cancer therapy. To date, approximately 20 clinical trials have been initiated using siRNA-based therapeutics. However, several barriers still exist to achieving effective and controlled *in vivo* delivery and these limits the use of siRNAs in the clinic. Post-intravenous injection, the siRNA complex must navigate the circulatory system of the body while avoiding kidney filtration, uptake by phagocytes, aggregation with serum proteins and enzymatic degradation by endogenous nucleases (33,34). The current siRNA delivery systems for cancer therapy mainly include chemical modifications of siRNA, lipid-based, polymer-based, and conjugate siRNA delivery systems, as well as co-delivery of siRNA and anticancer drugs, and inorganic nanoparticles (35). These modifications help to address the problems associated with naked siRNA delivery and effectively introduce the siRNA inside the target cells. In this study, two transfection reagents have been tested, DharmaFECT transfection reagent 1 and jetPEI. The former is a lipid-based formulation and the latter is a linear polyethylenimine (PEI) derivative. Both of these reagents effectively delivered the STAT6 siRNAs into the cells, as STAT6 expression was significantly knocked down in both cases. Nevertheless, jetPEI, unlike DharmaFECT, has been successfully tested in several animal studies and is known to form stable complexes with the nucleic acid, protecting it from degradation. Moreover, good manufacturing practice (GMP) grade *in vivo*-jetPEI is being used in several ongoing preclinical studies and phase I and II clinical trials. Thus, this makes jetPEI an excellent candidate for future animal and clinical studies using the STAT6 siRNA sequences used in this study.

In conclusion, all four STAT6 siRNA sequences significantly silenced STAT6 expression, reduced the number of live HT-29 cells and induced HT-29 apoptosis. Consequently, all four sequences, especially STAT6.1 and STAT6.4, are good candidates to develop as treatments of CRC. Animal studies using immunocompromised mice with human colon cancer xenografts are currently being planned. These will permit the determination of the *in vivo* effectiveness of the STAT6 siRNA sequences. The effectiveness of the STAT6 sequences in other cancers is also being tested. The experiments conducted in HT-29 cells are being reproduced in STAT6-expressing breast cancer cells and the results are promising.

## Supporting information

**S1Fig.**
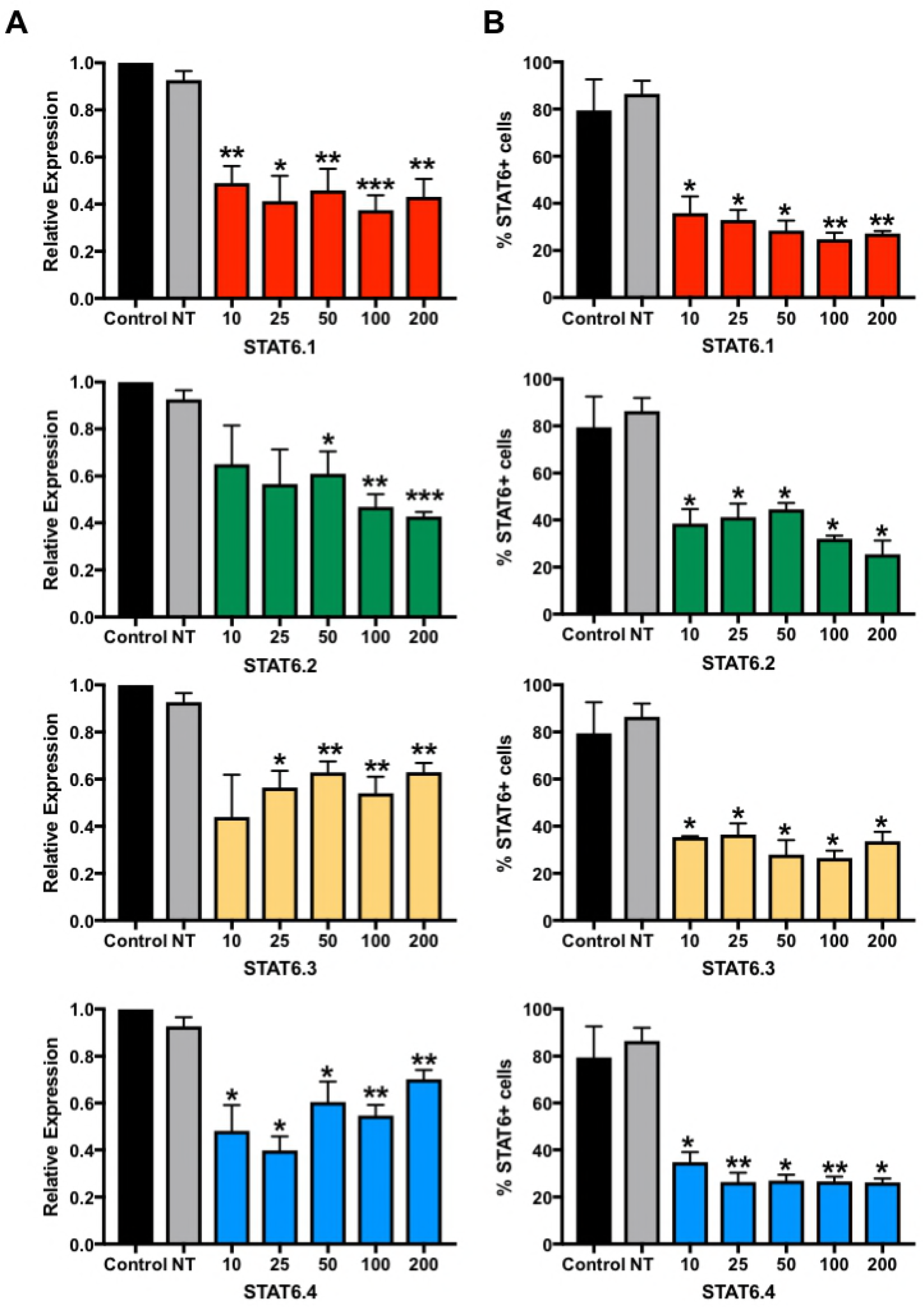
Optimal dose of STAT6 siRNA sequences. (A) STAT6 mRNA level measure. The graphs represent the mean ± SEM of 3 independent experiments. Values were obtained by real-time PCR and results were analysed by ∆∆Ct method for relative quantifications. The fold change is represented on the Y axis, and values are normalized to control cells. (B) STAT6 protein level analysis. The graphs represent the mean of the percentage of STAT6 positive cells ± SEM of 2 independent experiments obtained by flow cytometry. The percentage of STAT6 positive cells is represented on the Y axis. STAT6 siRNAs and non-targeting siRNA were used at 10, 25, 50, 100 and 200 nM as the final concentration. Control cells were non-transfected cells and STAT6 siRNA sequences 1, 2, 3 and 4 and non-targeting siRNA are denoted as STAT6.1, STAT6.2, STAT6.3 and STAT6.4 and NT, respectively.

**S2Fig.**
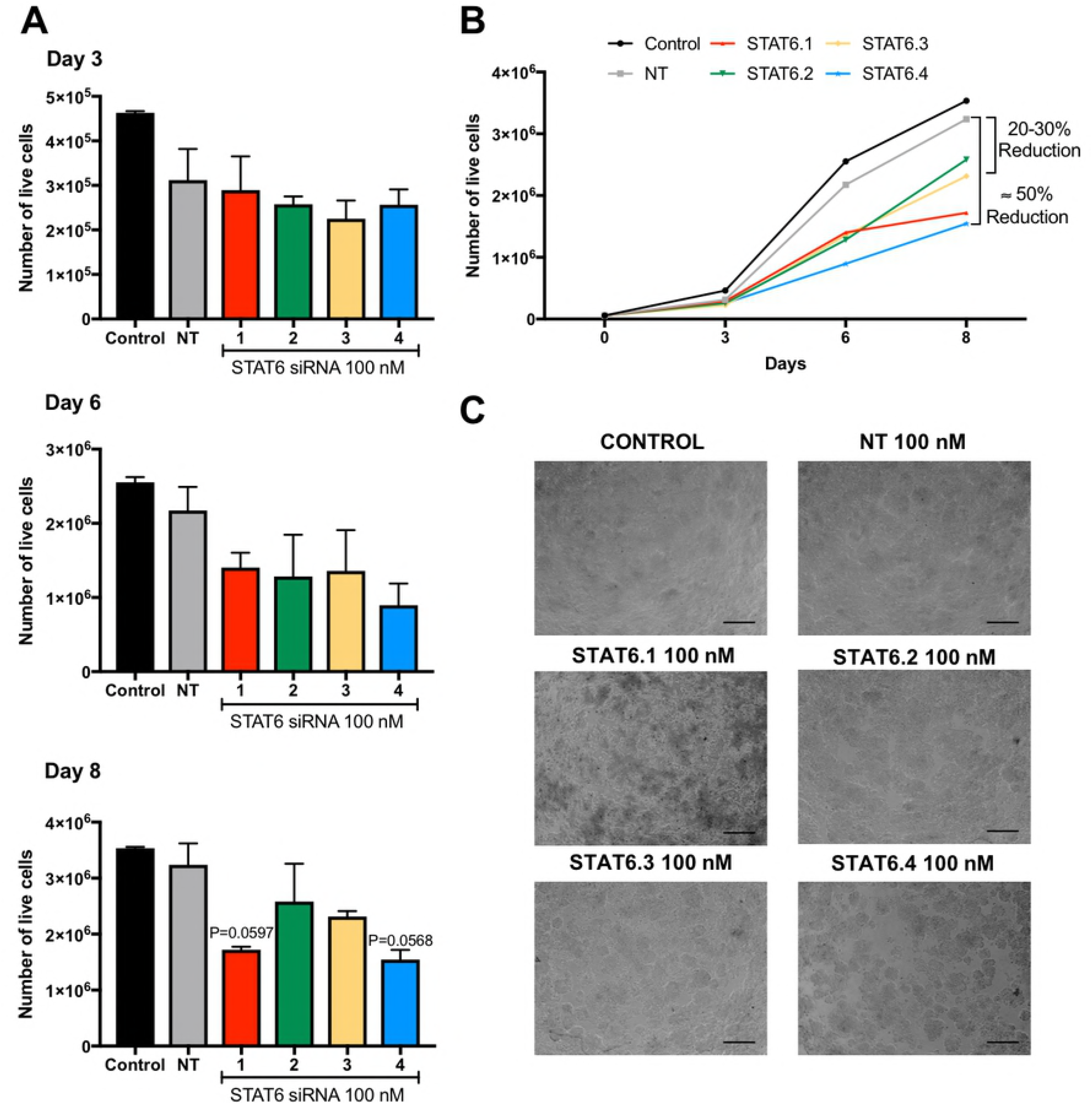
Cell proliferation using 100 nM STAT6 siRNA sequences 1 to 4. (A) Number of live cells measured at day 3, 6 and 8 of culture. The graphs represent the mean ± SEM of 2 independent experiments. (B) The graph shows how cells grew over time and represents the mean ± SEM of the independent experiments shown in A. The number of live cells was calculated as detailed in the material and methods section using NucleoCounter NC-100. STAT6 siRNAs and non-targeting siRNA were used at 100 nM as final concentration. (C) Inverted microscope image taken at day 8 of culture. Control cells were non-transfected cells and STAT6 siRNA sequences 1, 2, 3 and 4 and non-targeting siRNA are denoted as STAT6.1, STAT6.2, STAT6.3 and STAT6.4 and NT, respectively. The percentage of reduction of the number of live cells is calculated by comparison between the mean of NT *vs*. the mean of STAT6 siRNAs.

## References

1. Ferlay J, Soerjomataram I, Dikshit R, Eser S, Mathers C, Rebelo M, et al. Cancer incidence and mortality worldwide: Sources, methods and major patterns in GLOBOCAN 2012: Globocan 2012. Int J Cancer. 2015 Mar 1;136(5):E359–86.

2. Stewart BW, Wild C. International Agency for Research on Cancer, World Health Organization. World cancer report 2014 [Internet]. 2014 [cited 2018 Oct 8]. Available from: http://libweb.iaea.org/library/eBooks/World-Cancer-Report2014.pdf

3. Mármol I, Sánchez-de-Diego C, Pradilla Dieste A, Cerrada E, Rodriguez Yoldi M. Colorectal Carcinoma: A General Overview and Future Perspectives in Colorectal Cancer. Int J Mol Sci. 2017 Jan 19;18(1):197.

4. Darnell JE. STATs and Gene Regulation. Science. 1997 Sep 12;277(5332):1630.

5. Bowman T, Garcia R, Turkson J, Jove R. STATs in oncogenesis. Oncogene. 2000 May 22;19:2474.

6. Johnson DE, O’Keefe RA, Grandis JR. Targeting the IL-6/JAK/STAT3 signalling axis in cancer. Nat Rev Clin Oncol. 2018 Feb 6;15:234.

7. Hou J, Schindler U, Henzel W, Ho T, Brasseur M, McKnight S. An interleukin-4-induced transcription factor: IL-4 Stat. Science. 1994 Sep 16;265(5179):1701.

8. Kotanides H, Reich N. Requirement of tyrosine phosphorylation for rapid activation of a DNA binding factor by IL-4. Science. 1993 Nov 19;262(5137):1265.

9. Mikita T, Campbell D, Wu P, Williamson K, Schindler U. Requirements for interleukin-4-induced gene expression and functional characterization of Stat6. Mol Cell Biol. 1996 Oct;16(10):5811–20.

10. Wang C, Zhu C, Wei F, Zhang L, Mo X, Feng Y, et al. Constitutive Activation of Interleukin-13/STAT6 Contributes to Kaposi’s Sarcoma-Associated Herpesvirus-Related Primary Effusion Lymphoma Cell Proliferation and Survival. Longnecker RM, editor. J Virol. 2015 Oct 15;89(20):10416–26.

11. Li BH, Yang XZ, Li PD, Yuan Q, Liu XH, Yuan J, et al. IL-4/Stat6 activities correlate with apoptosis and metastasis in colon cancer cells. Biochem Biophys Res Commun. 2008 May 2;369(2):554–60.

12. Ostrand-Rosenberg S, Grusby MJ, Clements VK. Cutting Edge: STAT6-Deficient Mice Have Enhanced Tumor Immunity to Primary and Metastatic Mammary Carcinoma. J Immunol. 2000 Dec 1;165(11):6015–9.

13. Ostrand-Rosenberg S, Sinha P, Clements V, Dissanayake SI, Miller S, Davis C, et al. Signal transducer and activator of transcription 6 (Stat6) and CD1: inhibitors of immunosurveillance against primary tumors and metastatic disease. Cancer Immunol Immunother. 2004 Feb 1;53(2):86–91.

14. Uhlén M, Björling E, Agaton C, Szigyarto CA-K, Amini B, Andersen E, et al. A Human Protein Atlas for Normal and Cancer Tissues Based on Antibody Proteomics. Mol Cell Proteomics. 2005 Dec;4(12):1920–32.

15. Nappo G, Handle F, Santer FR, McNeill RV, Seed RI, Collins AT, et al. The immunosuppressive cytokine interleukin-4 increases the clonogenic potential of prostate stem-like cells by activation of STAT6 signalling. Oncogenesis. 2017 May 29;6(5):e342.

16. Gooch JL, Christy B, Yee D. STAT6 Mediates Interleukin-4 Growth Inhibition in Human Breast Cancer Cells. Neoplasia. 2002;4(4):324–31.

17. Zhang WJ, Li BH, Yang XZ, Li PD, Yuan Q, Liu XH, et al. IL-4-induced Stat6 activities affect apoptosis and gene expression in breast cancer cells. Cytokine. 2008 Apr;42(1):39–47.

18. Wang C-G. EZH2 and STAT6 expression profiles are correlated with colorectal cancer stage and prognosis. World J Gastroenterol. 2010;16(19):2421.

19. Street W. Colorectal Cancer Facts & Figures 2017-2019. :40.

20. Fire A, Xu S, Montgomery MK, Kostas SA, Driver SE, Mello CC. Potent and specific genetic interference by double-stranded RNA in Caenorhabditis elegans. Nature. 1998 Feb 19;391:806.

21. Agrawal N, Dasaradhi PVN, Mohmmed A, Malhotra P, Bhatnagar RK, Mukherjee SK. RNA Interference: Biology, Mechanism, and Applications. Microbiol Mol Biol Rev. 2003 Dec 1;67(4):657–85.

22. Carthew RW, Sontheimer EJ. Origins and Mechanisms of miRNAs and siRNAs. Cell. 2009 Feb;136(4):642–55.

23. Chakraborty C, Sharma AR, Sharma G, Doss CGP, Lee S-S. Therapeutic miRNA and siRNA: Moving from Bench to Clinic as Next Generation Medicine. Mol Ther - Nucleic Acids. 2017 Sep;8:132–43.

24. Walker W, Hopkin M. (54) MATERALS AND METHODS FOR TREATMENT OF ALLERGIC DISEASE. :46.

25. Merk BC, Owens JL, Lopes M-BS, Silva CM, Hussaini IM. STAT6 expression in glioblastoma promotes invasive growth. BMC Cancer [Internet]. 2011 Dec [cited 2018 Sep 4];11(1). Available from: http://bmccancer.biomedcentral.com/articles/10.1186/1471-2407-11-184

26. Cui X, Zhang L, Luo J, Rajasekaran A, Hazra S, Cacalano N, et al. Unphosphorylated STAT6 contributes to constitutive cyclooxygenase-2 expression in human non-small cell lung cancer. Oncogene. 2007 Jun;26(29):4253–60.

27. Wang N, Tao L, Zhong H, Zhao S, Yu Y, Yu B, et al. miR-135b inhibits tumour metastasis in prostate cancer by targeting STAT6. Oncol Lett. 2016 Jan;11(1):543–50.

28. Binnemars-Postma K, Bansal R, Storm G, Prakash J. Targeting the Stat6 pathway in tumor-associated macrophages reduces tumor growth and metastatic niche formation in breast cancer. FASEB J. 2018 Feb;32(2):969–78.

29. Yan D, Wang H-W, Bowman RL, Joyce JA. STAT3 and STAT6 Signaling Pathways Synergize to Promote Cathepsin Secretion from Macrophages via IRE1α Activation. Cell Rep. 2016 Sep;16(11):2914–27.

30. Cao H, Zhang J, Liu H, Wan L, Zhang H, Huang Q, et al. IL-13/STAT6 signaling plays a critical role in the epithelial-mesenchymal transition of colorectal cancer cells. Oncotarget [Internet]. 2016 Sep 20 [cited 2018 Sep 4];7(38). Available from: http://www.oncotarget.com/fulltext/11282

31. Zhang M, Zhou Y, Zhang W, Zhang X, Pan Q, Ji X, et al. Apoptosis induced by short hairpin RNA-mediated STAT6 gene silencing in human colon cancer cells. Chin Med J (Engl). 2006 May;119(10):801–8.

32. Rao DD, Vorhies JS, Senzer N, Nemunaitis J. siRNA vs. shRNA: Similarities and differences. Adv Drug Deliv Rev. 2009 Jul;61(9):746–59.

33. Alexis F, Pridgen E, Molnar LK, Farokhzad OC. Factors Affecting the Clearance and Biodistribution of Polymeric Nanoparticles. Mol Pharm. 2008 Aug;5(4):505–15.

34. Whitehead KA, Langer R, Anderson DG. Knocking down barriers: advances in siRNA delivery. Nat Rev Drug Discov. 2009 Feb 1;8:129.

35. Singh A, Trivedi P, Jain NK. Advances in siRNA delivery in cancer therapy. Artif Cells Nanomedicine Biotechnol. 2018 Feb 17;46(2):274–83.

